# Kalman-filter Force Inference: an estimation framework for cellular forces from temporal evolution of epithelial morphogenesis

**DOI:** 10.64898/2025.12.19.695339

**Authors:** Goshi Ogita, Takemasa Miyoshi, Tatsuo Shibata

## Abstract

Epithelial morphogenesis is orchestrated by cellular forces, such as cell junctional tension and cellular pressure. Therefore, elucidating the spatiotemporal distribution of these forces and their dynamics is paramount to understand morphogenesis during development. Over the past decade, various force inference methods have been proposed to estimate cellular forces based on the shape and geometry of cells within epithelial tissues. Most of these methods were developed under the assumption of static force balance, which neglects the cell deformation, thereby limiting their applicability to tissue undergoing dynamic deformation. To address this issue, we develop a novel method to accurately infer cellular forces from time-lapse imaging data of epithelial deformation, without relying on the static force balance assumption. Our method is based on two fundamental assumptions. First, the forces exerted by cells are balanced with dissipative forces (such as viscous and frictional forces) at each vertex arising from cell deformation, referred to as the dynamic force balance. Second, cellular forces vary smoothly over time. By formalizing these assumptions using a Bayesian framework, we construct a new approach of force inference, named Kalman-filter Force Inference (KFI). The effectiveness of our method is evaluated using synthetic data of a deforming tissue generated by numerical simulations of the cell vertex model. The results demonstrate the accurate estimation of cellular force dynamics. Furthermore, our systematic evaluation reveals the capability of our method to accurately estimate cellular forces across tissues with diverse mechanical parameters. Finally, we assessed the robustness of our method to observation noise. We anticipate that Kalman-filter Force Inference will broaden the applicability of force inference techniques and significantly contribute to our understanding of epithelial mechanics.

## 1 Introduction

During organismal development, living tissues undergo pronounced shape changes to form complex and functional organs and body structures. Epithelium is the most common tissue that shapes the organs. It consists of tightly adhered cells that line both external and internal surfaces. The deformation of the epithelial tissue is directed by mechanical processes. Therefore, how such mechanical processes are produced and regulated is a major focus in developmental biology.

In epithelial tissues, actomyosin cortices localize at cell-cell junctions, forming tensile networks that are thought to govern epithelial mechanics. Epithelial morphogenesis is viewed as a mechanical process of such a network, characterized by junctional tension and intercellular pressure within individual cells [Fig. 1(a) left and middle] [1]. These cellular forces collectively drive cell deformation and rearrangement, ultimately leading to the tissue-scale deformation. The overall tissue property as a material, such as viscoelasticity, also plays an important role in tissue-scale deformation, [2, 3, 4]. The tissue material properties are regulated by the cellular mechanical parameters such as junction contractility, cell-cell adhesion strength, and cell compression modulus [2, 5, 6]. In this context, quantifying how cellular forces are spatiotemporally distributed is the first step to elucidating epithelial morphogenesis.

**Figure 1:**
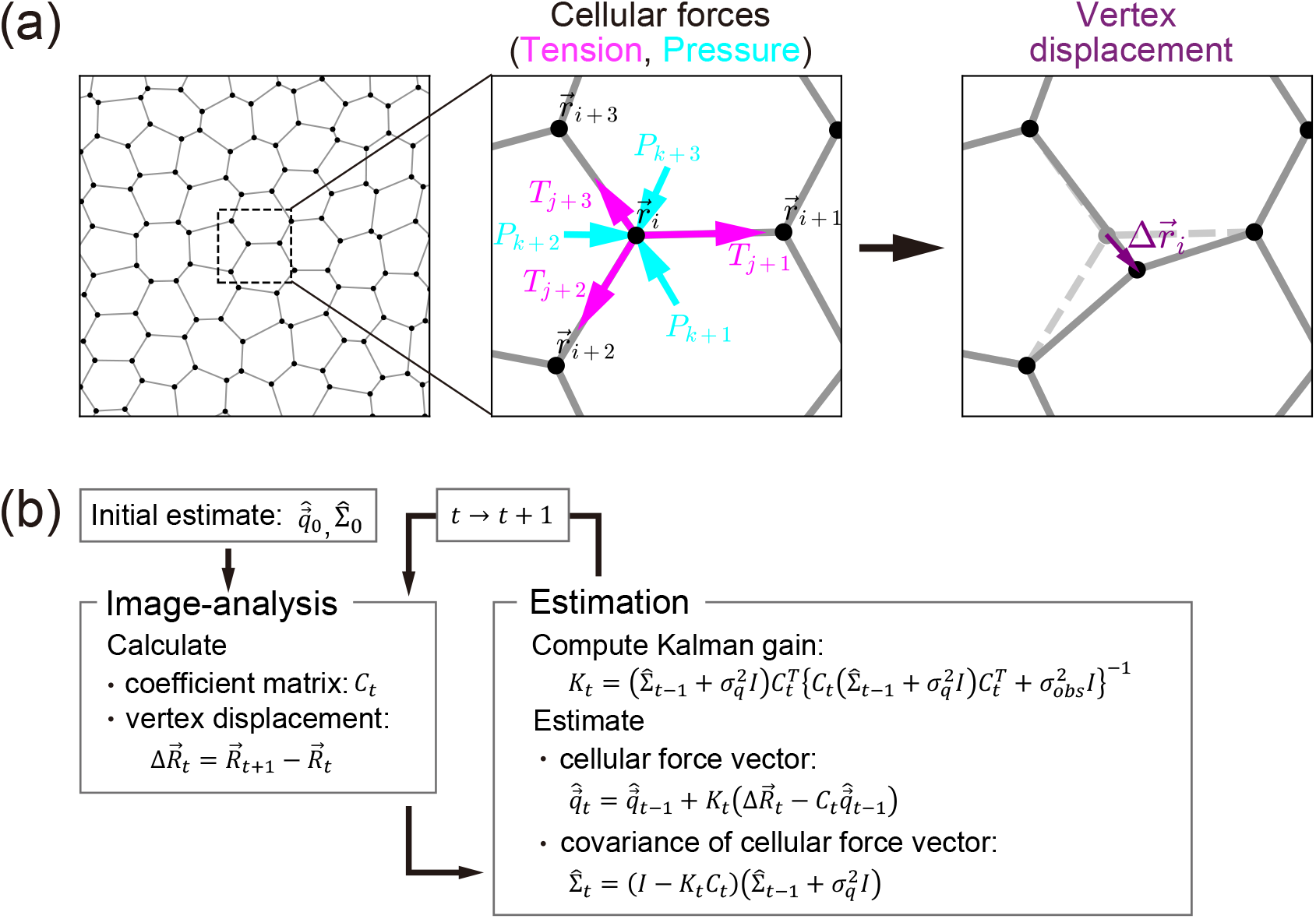
Kalman-filter Force Inference (a) The dynamic force balance between cellular forces and dissipative forces in the epithelial tissue. Epithelial cells, depicted as polygons, exhibit tight adhesion at their junctions (edges) (left panel). Tension (magenta arrows) acts along the junctions, while pressure (cyan arrows) is generated within the cells (middle panel). The net forces of junctional tension and cellular pressure displace cell vertices (black dots) and are balanced by the resulting dissipative forces arising from cell deformation (right panel). (b) Estimation cycle of Kalman-filter Force Inference (KFI); see section 2.1 for details.

Several methods for measuring cellular forces have been applied to tissues in various organisms [7, 8, 9]. Among these, force inference methods have been widely used to estimate cellular forces from the geometry of cell shapes and arrangements obtained from microscopy images [10, 11]. These methods are non-invasive, easy to apply, and high-throughput, and have therefore been applied to a wide range of tissues, contributing to our understanding of the roles of cellular forces in epithelial morphogenesis and the maintenance of tissue homeostasis [12, 13, 14, 15, 16, 17, 18, 19]. Various force inference methods have been proposed since the 2010s. These methods share a common strategy of solving force balance equations—mathematical expressions that relate cellular forces (junctional tensions and cellular pressures) to cell geometries, either at mechanical equilibrium or during tissue deformation.

The first force inference method, termed Video Force Microscopy (VFM), was proposed in 2010 to estimate cellular forces during ventral furrow formation in Drosophila embryos [20]. In this method, cross-sections of Drosophila embryos are analyzed, with individual cells represented as quadrilaterals, and force balance equations are formulated between junctional tensions, cellular pressures, and dissipative forces such as friction and viscous forces arising during cell deformation. Because the resulting system of equations is underdetermined, VFM exploits temporal information from time-lapse data and employs recursive least squares with a forgetting factor to solve the equations and estimate the cellular forces.

Cellular forces on the apical surface of epithelial sheets were first estimated by methods called Mechanical Stress Inference (MSI) [12] and Bayesian Force Inference (BFI) [15], both proposed in 2012. In these methods, cell shapes are approximated by polygons, and static force balance is considered among the cellular forces without accounting for contributions from tissue deformation. Additional constraints are imposed on the mean value of junctional tensions, such as the assumption that the average tension is positive. To solve the resulting underdetermined equations, MSI uses the Moore–Penrose pseudoinverse, whereas BFI adopts a Bayesian statistical framework with a likelihood function that embeds the static force balance equations and a prior distribution constraining the mean junctional tension, both modeled as normal distributions. The maximum a posteriori (MAP) estimate of the forces is then obtained as the force inference result.

Force balance equations can also be solved by taking into account the curvature of cell junctions. The Cellular Force Inference Toolkit (CellFIT), proposed in 2014, combines force balance equations for junctional tensions at cell vertices with the Young–Laplace equation between junctional tension, cellular pressure, and junction curvature along cell-cell junctions, thereby rendering the problem overdetermined and enabling force estimation without additional constraints [21]. Temporal information was subsequently incorporated, within a framework similar to CellFIT, in Dynamic Local Intercellular Tension Estimation (DLITE), proposed in 2019, which improves estimation stability by using the force estimate at the previous time point as the initial value for the numerical solution of the static force balance equation [22]. The Variational Method of Stress Inference (VMSI), proposed in 2020, achieved more robust estimation by further incorporating geometric cell information, such as junction curvature—previously treated only as observables—as additional parameters to be inferred [14]. Furthermore, ForSys, proposed in 2024, extends CellFIT to cellular force dynamics by incorporating dissipative forces, as in VFM [23].

Most methods focus on static force balance, in which contributions from cell and tissue deformations are not considered, except for VFM and ForSys. This assumption is reasonable when tissue deformation is slow or almost absent. It has been reported that cellular forces estimated by a static force inference method deviate from forces inferred from myosin distributions [14]. This discrepancy can be attributed to the neglect of dissipative forces arising from cell and tissue deformations, motivating the development of stable dynamic force inference methods that explicitly incorporate these deformations.

Existing methods such as MSI and BFI often impose constraints on junctional tension, such as tension positivity, to overcome the underdetermined nature of the equations. While such constraints improve robustness to observational errors and missing data [15, 24, 25], they can become problematic when the assumed tension properties are not satisfied. Recent studies have shown that estimation accuracy decreases when tissue material properties become more fluid-like [26], likely because such tissues may exhibit negative junctional tensions that violate the positivity constraint. This motivates the development of methods that relax constraints on junctional tension.

Therefore, in the present paper, we develop a novel dynamic force inference method within a Bayesian statistical framework. In this method, we consider dynamic force balance equations that explicitly include contributions from cell deformation as dissipative forces. In line with BFI, the likelihood function embeds the dynamic force balance equations. The prior distribution imposes temporal continuity on cellular forces, similar to the approach used in VFM and DLITE [20, 22]. This assumption is inspired by Bayesian approaches used to quantify in-plane stresses of cultured epithelial monolayer from traction force microscopy data, in particular by the extension from Bayesian Inversion Stress Microscopy (BISM) to Kalman Inversion Stress Microscopy (KISM) [27, 28]. By combining these elements, we obtain a force inference scheme analogous to the Kalman filter, which is widely used in time-series analysis [29], and we term it Kalman-filter Force Inference (KFI). We evaluated the performance of KFI using synthetic datasets generated by numerical simulations, which encompassed a range of cellular force dynamics and tissue material properties. KFI accurately estimated cellular forces across a wide range of conditions and maintained reasonable accuracy even in the presence of observation noise. Additionally, the Kalman filter framework quantifies the uncertainty of each force estimate, which is one of its key advantages. With these advantages, KFI has the potential to broaden the applicability of force inference methods and to uncover new roles of dynamical forces in epithelial morphogenesis.

## 2 Results

### 2.1 Kalman-filter Force Inference

In this section, we present Kalman-filter Force Inference (KFI), a novel method for estimating cellular forces from time-lapse imaging data of epithelial tissue deformation. We first formulate the dynamic force balance equations that govern epithelial deformation (Section 2.1.1), which provide the foundation for two complementary derivations of KFI (Sections 2.1.2 and 2.1.3). We then derive KFI using Bayes’ theorem by treating the dynamic force balance equations as a probabilistic inference problem, following the approach of Bayesian force inference [15] (Section 2.1.2). Finally, we show that KFI can be equivalently derived within the Kalman filter framework by representing tissue dynamics as a state-space model (Section 2.1.3). Both approaches yield identical estimation equations, demonstrating the mathematical equivalence between the Bayesian inference framework and the Kalman filtering approach.

#### 2.1.1 Dynamic force balance equation for epithelial deformation

To develop a time-dependent force inference method, we first consider the force balance between the cellular forces, such as junctional tension and cellular pressure, and the dissipative forces arising from the cellular motions, including cell deformation. This relationship is described by a dynamic force balance equation or, equivalently, an equation of motion.

The epithelial tissue is approximately described by a polygonal tesselation. Individual epithelial cell shape is given by polygons with vertices and edges. Vertex *i* at position 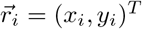, as shown in Fig. 1(a), experiences the tension *T*_*j*_ of cell-cell junction *j* and the pressure *P*_*k*_ of cell *k*. The total cellular force at vertex 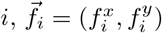, is then given by

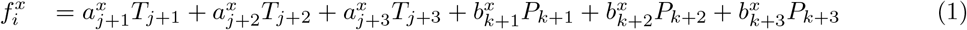

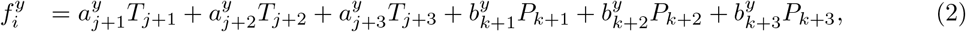

where the coefficients 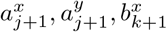 and 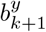 can be calculated from the coordinates of the cell vertices [15]. For example, 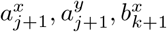 and 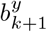 are given by 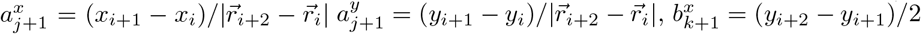 and 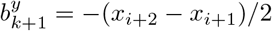.

We also consider frictional forces, which may arise from vertex displacements or cell-cell junction deformations [30, 31, 32]. Here, our purpose is not to investigate the effect of a specific form of frictional force on tissue mechanics. Instead, we adopt a general form of such frictional forces, which can be written as

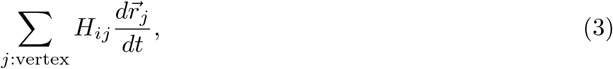

where *H* is a symmetric 2*N*_*V*_ × 2*N*_*V*_ friction matrix [32].

We neglect inertial effects as is standard for tissue dynamics [33, 34]. Then, the dynamic force balance equation is given by

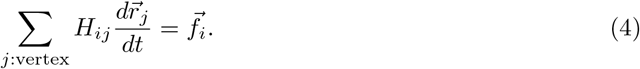

In the previous static force inference methods, a static force balance relation 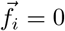 was assumed, while the present method considers a dynamic force balance relation given by Eq.(4).

When the epithelial tissue consists of *N*_*C*_ cells with *N*_*E*_ cell-cell junctions and *N*_*V*_ cell vertices, a combination of all tensions and pressures gives a (*N*_*E*_ + *N*_*C*_)-dimensional vector 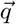 given by 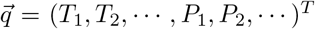. Then, a 2*N*_*V*_-dimensional vector 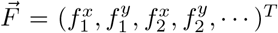, which consists of forces of *N*_*V*_ cell vertices, is given by a linear combination of tensions and pressures as

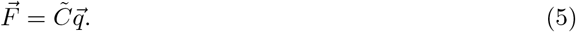

where 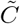 is a 2*N*_*V*_ × (*N*_*E*_ + *N*_*C*_) matrix composed of the coefficients such as 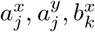, and 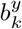. Combining the coordinates of the vertices as 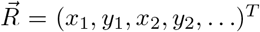, the dynamic force balance equation for the tissue given by Eq.(4) can be collectively written as

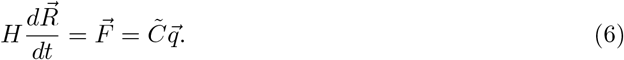

With a finite-time discretization of Eq.(6) up to the order of Δ*t*, the displacement of cell vertices 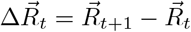 can be written as

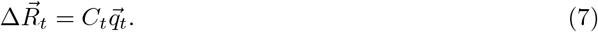

with 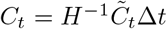. Here, both 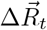 and *C*_*t*_ can be constructed from the information extracted from the image data at times *t* + 1 and *t*. Therefore, by solving this equation with respect to 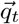, we can estimate the cellular forces such as junctional tension *T*_*j*_ and cellular pressure *P*_*k*_ at time *t*. However, this equation cannot be solved independently, since the number of unknown variables exceeds the number of available conditions [15, 12].

#### 2.1.2 Force inference method formulated within a Bayesian statistical framework

Here, we adopt Bayes’ theorem to resolve this indeterminacy as in the case of Bayesian force inference [15]. According to Bayes’ theorem, the prior distribution 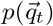, the likelihood function 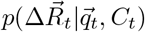, and the posterior distribution 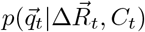 are related by the following proportionality:

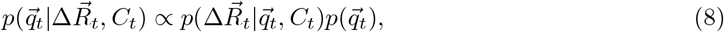

In the present formulation, we suppose that 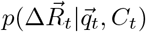 and 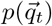 are well-described by Gaussian function, giving 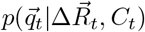 to be Gaussian.

We first consider the likelihood function based on Eq. (7). When determining the positions of cell vertices, there is always uncertainty due to insufficient microscopic resolution, image pixelization, and segmentation, which can be considered as observation noises. The observation noise can be modeled by an additive noise 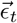 to Eq. (7) with 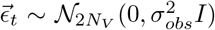, giving the following observation equation:

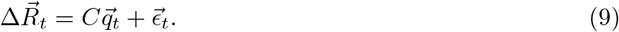

Thus, we obtain the likelihood function as

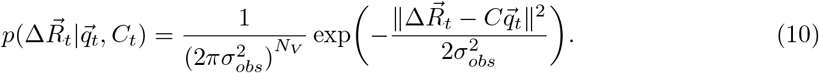

For the prior distribution, reflecting our lack of knowledge about the time evolution of the cellular forces, we suppose that the cellular forces fluctuate around their value at the previous time step as

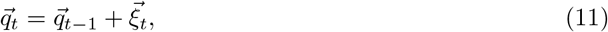

where 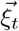 is a Gaussian noise with zero mean and covariance 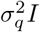. Thus, for the posterior distribution of 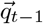 with mean 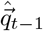 and covariance 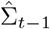, the prior distribution 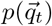 follows 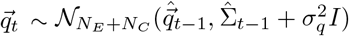, that is

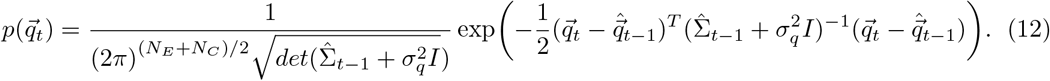

With the likelihood function Eq. (10) and the prior distribution Eq.(12), from Bayes’ theorem Eq.(8), the mean and the covariance of the posterior distribution, 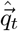 and 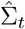 are respectively given by

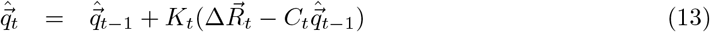

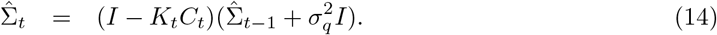

Here *K*_*t*_ is called the Kalman gain in the formulation of the Kalman filter [29], given by

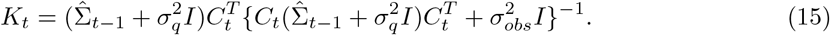

The posterior distribution of 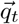 at *t* is used as the prior distribution for 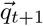 at *t* + 1, giving an analytical cycle. Since this analytical cycle is equivalent to that of the Kalman filter, we have named our method Kalman-filter Force Inference (KFI).

To implement this recursive estimation for a time-lapse movie recorded over *T* + 1 timesteps (from *t* = 0 to *t* = *T*), we initialize the process at time *t* = 0 by setting the mean 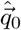 and covariance matrix 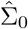 of the cellular forces. A time-lapse movie, recorded over *T* + 1 timesteps (from *t* = 0 to *t* = *T*), provides the time-series of vertex coordinates 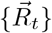. From the vertex coordinates, we calculate the vertex displacements 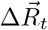 and construct the observation matrices *C*_*t*_ based on tissue geometry at each time step. We then estimate *T* sets of cellular force vector 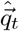 and their variance matrix 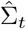 for *t* = 1, 2, …, *T*. 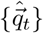 represents the time-series estimates of cellular forces. Specifically, the *j*-th component of 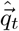 corresponds to 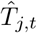, which is the estimated tension of the *j*-th cell-cell junction at time *t*. Similarly, the (*N*_*E*_ + *k*)-th component of 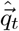 represents 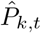, which is the estimated pressure of the *k*-th cell at time *t*. 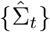 quantifies the uncertainty in these cellular force estimates. The (*j, j*) component of 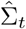 corresponds to the variance of the tension for the *j*-th cell-cell junction. Likewise, the (*N*_*E*_ + *k, N*_*E*_ + *k*) component of 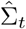 represents the variance of the pressure for the *k*-th cell. Using these variance estimates, the 95% highest posterior density interval (HDI) for 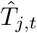 is calculated, for example, as 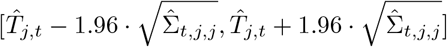.

#### 2.1.3 Force inference method formulated within a Kalman-filter framework using a state-space model

In the previous subsection, we constructed KFI using a Bayesian approach based on Bayes’ theorem. Here, we show that KFI can equivalently be formulated by representing epithelial tissue dynamics as a state-space model and applying the Kalman filter. A state-space model consists of two components: a system model describing the dynamics of the state variables and an observation model relating these states to observable quantities. In our formulation, cellular forces are treated as the state variables, whereas vertex positions and their displacements are treated as observable quantities. The Kalman filter then provides an optimal recursive solution for estimating the state variables from these observations.

The system model describes the temporal evolution of cellular forces. In the absence of detailed knowledge about force dynamics, we adopt the simplest model, a random walk, in which cellular forces fluctuate around their previous values. This is expressed by Eq. (11), where the state vector 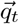 represents the cellular forces at time *t*, and 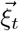 represents the system noise with zero mean and covariance 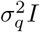. The system matrix is simply the identity matrix *I*, indicating that forces are expected to remain similar across consecutive time steps, subject to random perturbations. The observation model links the cellular forces to measurable vertex displacements. As derived from the dynamic force balance equation, the relationship between cellular forces and vertex displacement is given by Eq. (9), where 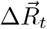 is the observation vector, *C*_*t*_ is the observation matrix constructed from tissue geometry, and 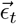 represents observation noise with zero mean and covariance 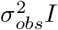 due to imaging limitations.

With this state-space representation, the Kalman filter provides a recursive algorithm to estimate cellular forces. The filtering process consists of two steps, namely prediction and update step, which are iterated sequentially for each time point. At time *t* = 0, the mean 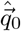 and covariance matrix 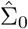 of the cellular forces are initialized. At each subsequent time step, the prediction step uses the posterior distribution from the previous time point to predict the prior distribution at time *t* based on the system model (Eq. 11). Because the cellular forces evolve as a random walk, the predicted mean remains 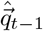, while the predicted covariance becomes 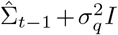, where the added term 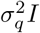 accounts for the stochastic fluctuations in force dynamics. The update step incorporates the newly observed vertex displacement 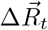 to refine the prediction. Using the observation model (Eq. 9) and Bayes’ theorem, the predicted distribution is updated to yield the posterior distribution with mean 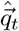 and covariance 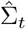, as given by Eqs. (13) and (14). In Eq. (13), the Kalman gain *K*_*t*_ given by Eq. 15 optimally weights the contribution of the new observation relative to the prediction.

By iterating the prediction and update steps from *t* = 1 to *t* = *T*, we obtain time-series estimates of the cellular forces 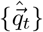 and their associated uncertainties 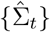 over all time points. This state-space formulation with Kalman filtering is mathematically equivalent to the Bayesian formulation presented in the previous subsection, demonstrating that KFI can be derived from two complementary perspectives.

### 2.2 Kalman-filter Force Inference estimated dynamic cellular forces with high accuracy

One advantage of Kalman-filter Force Inference (KFI) is that it explicitly considers the dynamic force balance described by Eq. (4), in contrast to the previous methods that assumed only static force balance between junctional tensions and cellular pressures. Consequently, KFI is expected to yield accurate estimations of the cellular forces in situations away from static force balance. To evaluate the performance of KFI under such conditions, we applied it to synthetic data of deforming epithelial tissue generated by numerical simulations of the cell vertex model, a widely used theoretical framework for modeling epithelial deformation (see 4.3 for details) [35, 36, 30, 31].

To drive the system away from static force balance, we introduced time-dependent sinusoidal forcing into the junctional tension in the cell vertex model [36, 37] as

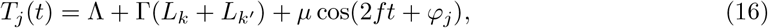

where Λ and Γ are the line tension and the cortical elasticity, respectively, determining the cellular mechanical parameters. *L*_*k*_ and *L*_*k*_^*′*^ are respectively the perimeter length of cells *k* and *k*^*′*^, which are separated by junction *j. µ, f*, and *φ*_*j*_ are the amplitude, frequency, and initial phase, respectively, controlling the sinusoidal forcing. Throughout this subsection, we set (Λ, Γ) = (0.1, 0.04). We incorporated the dissipative force arising from vertex displacement into the data generation and applied it consistently in the force inference method [*H* = *ηI* in Eq. 4].

Fig. 2 shows an example of synthetic data of square epithelial tissue with the parameter value (*µ, f*) = (0.1, 0.2), and the corresponding force inference using KFI. The snapshot images in Figs. 2(a) and (b) show the spatial distribution of junctional tensions and cellular pressures at *t* = 30.0 in the synthetic data [Fig. 2(a)] and the force inference data by KFI [Fig. 2(b)]. The images of synthetic and inferred data show good agreement, indicating that KFI estimated the cellular forces accurately. To see the agreement more quantitatively, we made a scatter plot between true and estimated values [Fig. 2(c)]. Most points were distributed along the diagonal line, with the correlation coefficients of 1.0 for tension and 0.99 for pressure, confirming the accuracy of the proposed method. The outliers that deviate from the diagonal line originate at the tissue boundary. The correlation coefficients at all time points were distributed ranging from 0.96 to 1.0 for tension and nearly 1 for pressure [Fig. 2(d)]. Actually, the correlation coefficients showed a high value immediately after the initial time steps. In fact, the estimated tension and pressure values at the particular cell-cell junction and cell quickly converged to their true values, respectively, immediately after the initial time point [Fig. 2(e)]. In addition to the force estimates, we plotted the 95% highest density interval (HDI; gray shade) as a measure of uncertainty, within which the tension value falls with 95% probability [38]. The 95% HDI was narrow relative to the amplitude of tension oscillation, indicating that KFI successfully estimated cellular forces with high accuracy throughout the time course.

**Figure 2:**
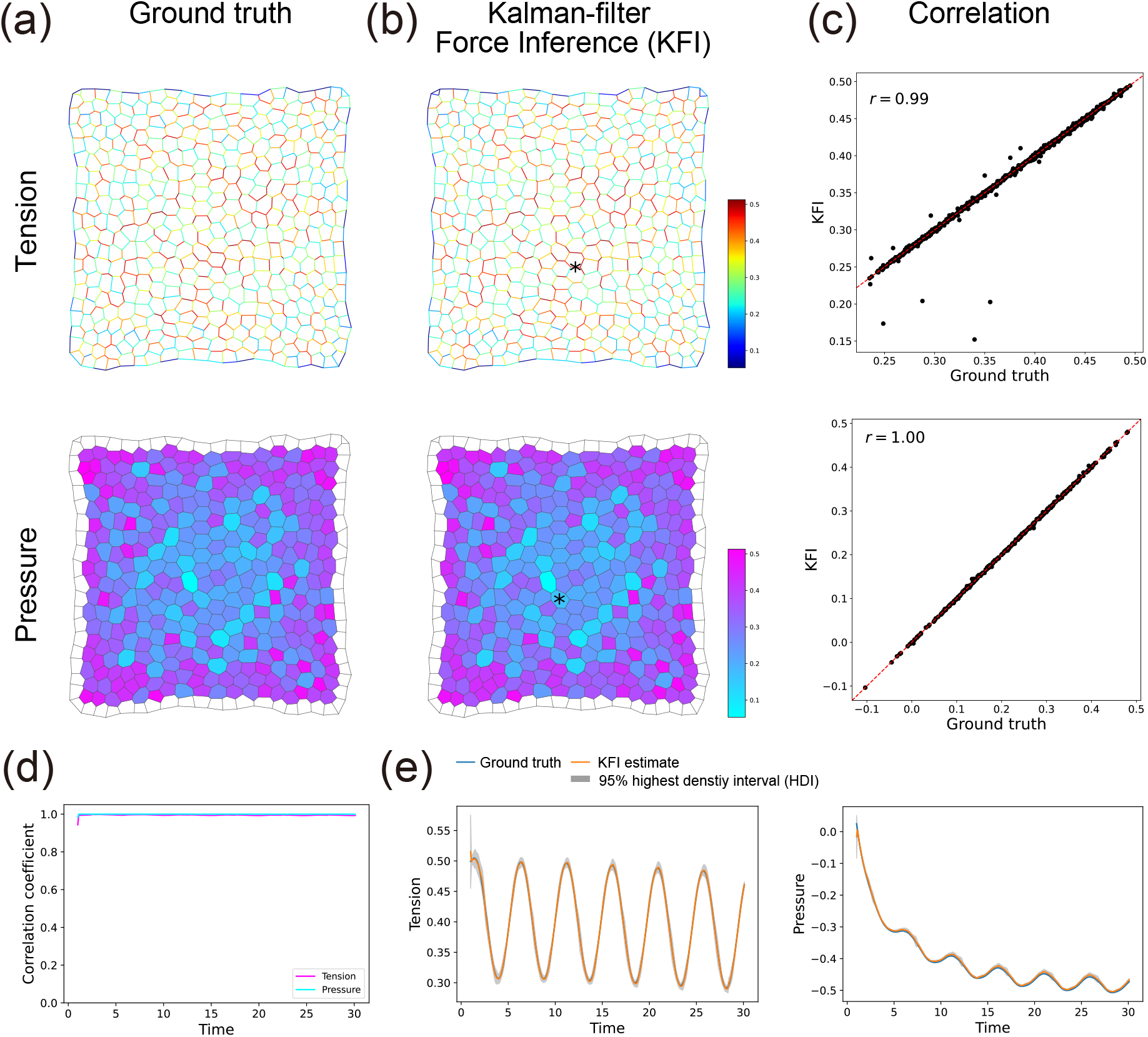
Application examples of Kalman-filter Force Inference. (a, b) The spatial distribution of the ground truth and the KFI estimates of the cellular force at *t* = 30.0. The top and bottom rows correspond to cell junctional tensions and cellular pressures, respectively. The magnitude of cellular forces is indicated by the colors exemplified in the color bar. (c) The correlation between the ground truth and the KFI estimates of the cellular forces at *t* = 30.0. The top and bottom panels correspond to cell junctional tensions and cellular pressures, respectively. The dashed line indicates the *y* = *x* line. The number in the upper left corner of each panel represents the Pearson correlation coefficient between the ground truth and the KFI estimates. (d) Correlation coefficients between the ground truth and the KFI estimates over the entire time course. The magenta and cyan lines correspond to the correlation coefficients for junction tension and cellular pressure, respectively. (e) Temporal dynamics of the ground truth and the KFI estimates of cellular forces (left: junctional tension; right: cellular pressure). The ground truth (blue line) and the estimate (orange line) are plotted against time. The gray shaded area indicates the 95% highest density interval of the estimates. The corresponding junction and cell are indicated by the asterisk in the top and bottom panels of (b), respectively.

Since previous methods assumed static force balance between junctional tensions and cellular pressures [10], their accuracy in estimating forces from the synthetic data used above is not expected to be as high as that of our proposed method. For comparative evaluation, we performed force inference on the same dataset using a conventional method. We selected Bayesian Force Inference (BFI) as the benchmark [see section 4.2 for details] [15], given that it shares the polygonal cell approximation assumption with KFI. As shown in Fig. S1(a), BFI estimates the junctional tensions and cellular pressures reflecting their spatial distribution at each time point. The correlations between the true and estimated values were 0.57 for tension and 0.62 for pressure for a particular time point, which were not as high as those of KFI [Fig. S1(b)]. For all time points, the correlation coefficients were distributed ranging from 0.4 to 0.6 for tension and 0.6 to 0.8 for pressure [Fig. S1(c)], suggesting that the estimation accuracy of BFI was lower than that of KFI.

We next study how the estimation accuracy depends on the deviation from the static force balance condition. The deviation can be controlled by the sinusoidal forcing in Eq. (16), parameterized by *µ* and *f*. In Fig. 3(a) and (d), we plotted the temporal average of the correlation coefficient between true and estimated cellular forces ⟨*r*⟩ up to *t* = 30.0 for different value of (*µ, f*). The value of ⟨*r*⟩ was distributed in the range 0.87 ~ 1.0 for tension and close to 1.0 for pressure, suggesting that the estimation accuracy of KFI was quite high in a wide range of *µ* and *f*. As shown in Fig. 3(b), the estimation accuracy of the junctional tension increased gradually with *µ*, suggesting that the dynamical change in the vertex position facilitates the junctional tension estimation. In contrast, the estimation accuracy of the junctional tension remained constant independent of *f*, suggesting the robustness of the method [Fig. 3(c)]. For the cellular pressure, the estimation accuracy remained high regardless of *µ* and *f* [Figs. 3(e),(f)].

**Figure 3:**
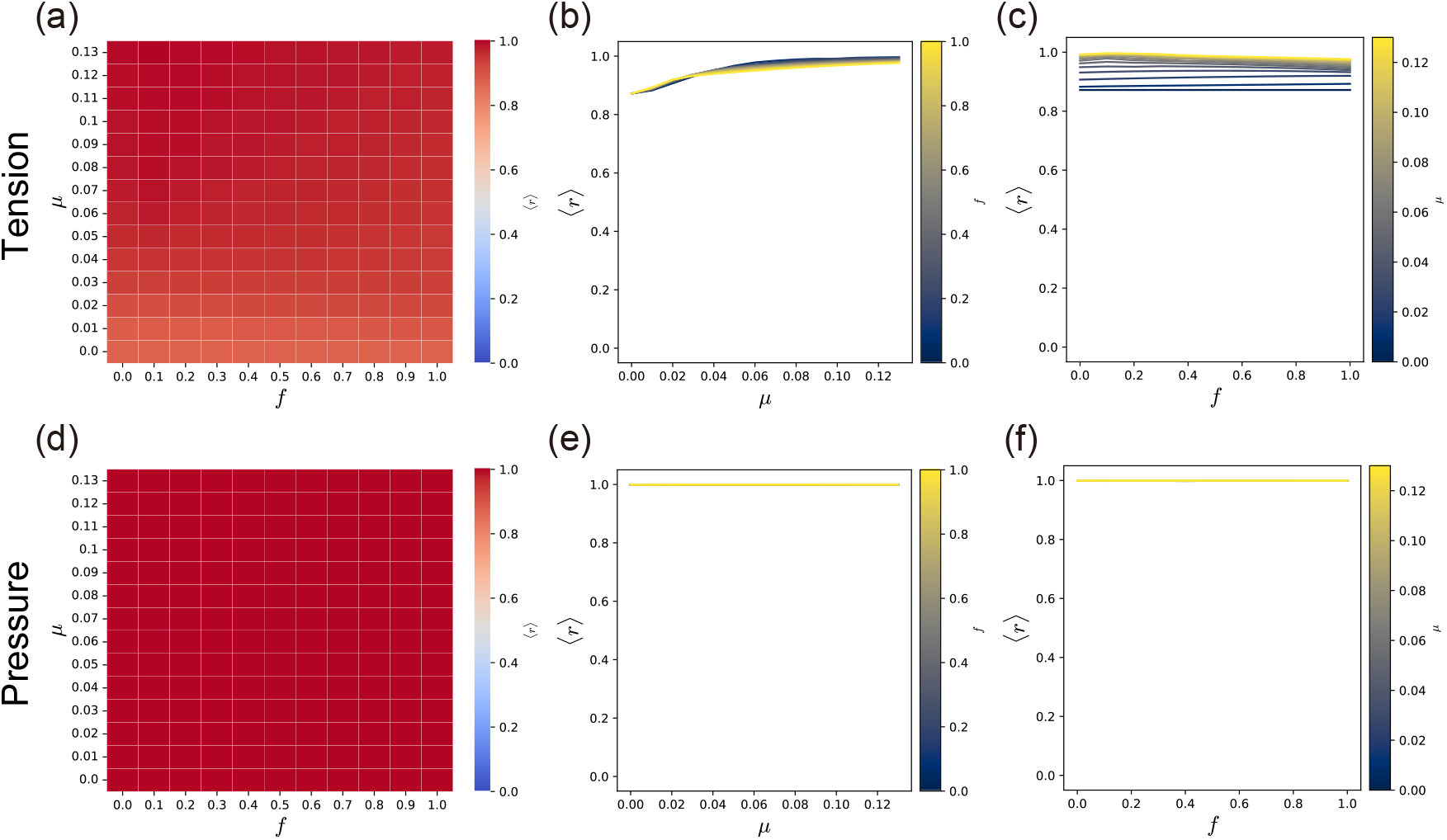
Estimation accuracy of Kalman-filter Force Inference (KFI) for various dynamic parameters (*µ, f*). (a-f) Parameter (*µ, f*) dependence of KFI’s estimation accuracy for junction tension (a-c) and cellular pressure (d-f). The mechanical parameters (Λ, Γ) were set to (0.1, 0.04). Heatmaps of estimation accuracy in the (*µ, f*) parameter space (junctional tension: a; cellular pressure: d). Colors in each cell represent the time average of the correlation coefficient between the ground truth and the estimation up to *t* = 30.0. Dependence of estimation accuracy on *µ* (junctional tension: b; cellular pressure: e) and *f* (junctional tension: c; cellular pressure: f).

We finally studied the estimation accuracy of BFI and how it depends on *µ* and *f* [Fig. S2]. As shown in Fig. S2 (a) and (d), the temporal average of the correlation coefficients ⟨*r*⟩ were distributed ranging from 0 to 0.70 for the tension and from 0.60 to 0.68 for the pressure, indicating that the estimation accuracy of BFI was lower than that of KFI. In particular, the estimation accuracy of the tension decreased as *f* increased [Fig. S2 (c)], suggesting that reliable force inference becomes difficult when vertex positions change rapidly and deviate from static force balance.

Overall, the results indicate that KFI provides accurate estimations of cellular forces in the situation far from static force balance.

### 2.3 Kalman-filter Force Inference accurately estimated cellular forces in tissues with a wide range of cellular mechanical parameters

Force inference methods should accurately estimate cellular forces regardless of the tissue material properties and cellular mechanical parameters. In cell vertex models, the tissue material property can be controlled by non-dimensional cellular mechanical parameters, such as 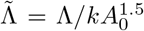 and 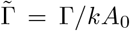. The parameter space of 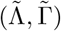 is separated into two regions based on its ground state. One is the hexagonal regime, where the ground state is a regular honeycomb network. The other is the soft-network regime, where the ground states are irregular soft networks and multiple configurations coexist [36, 39]. A recent report showed that the accuracy of BFI is high at the hexagonal regime, but declines at the soft-network regime [26]. This decline likely stems from the prior distribution of BFI, in which junctional tensions are assumed to be positive. This assumption becomes inappropriate for estimating cellular forces in the soft-network regime, where 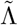 is negative and tensions can thus be negative. To quantify this, we calculated the time-averaged ratio of the mean to the standard deviation of junctional tension, ⟨*µ*_*T*_ */σ*_*T*_⟩. When ⟨*µ*_*T*_ */σ*_*T*_⟩*<* 1, the standard deviation exceeds the mean, indicating that junctional tensions frequently take negative values [Fig. S3(a)]. In the soft-network regime, ⟨*µ*_*T*_ */σ*_*T*_⟩ was indeed less than 1, confirming that negative tensions occur and contradict BFI’s assumption [Fig. S3(b)]. On the other hand, the prior distribution of KFI is defined by the posterior from the previous time point found in Eq. 12, allowing the Kalman filter to iteratively refine an inaccurate prior into an appropriate distribution, which results in the accurate estimation in the soft-network regime.

Here, we investigate how the accuracy of Kalman-filter Force Inference (KFI) depends on the tissue state. For various values of 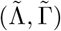, the time-averaged correlation coefficient ⟨*r*⟩ between the true and inferred values remained near 1 [Fig. 4], indicating that KFI works independently of the tissue state. In contrast, the BFI estimate was less accurate. The correlation coefficients for both tension and pressure were distributed from 0 to 0.6, dependent on 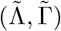. Notably, the estimation accuracy of BFI was significantly decreased in soft-network regime with low 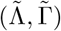, consistent with the previous study [26].

**Figure 4:**
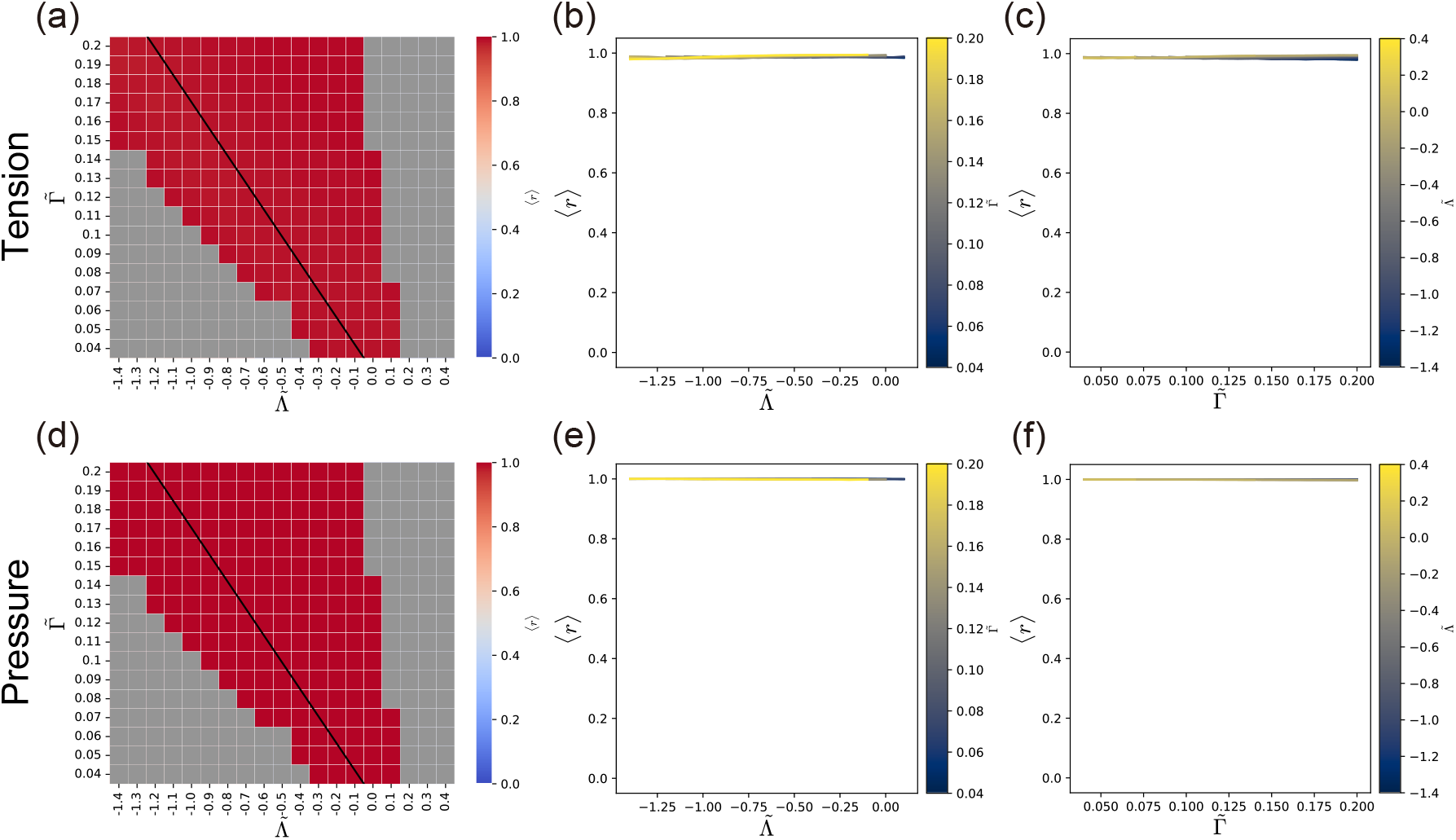
Estimation accuracy of Kalman-filter Force Inference (KFI) for various mechanical parameter 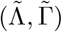. (a-f) Parameter 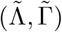 dependence of KFI’s estimation accuracy for junctional tension (a-c) and cellular pressure (d-f). The dynamics parameters (*µ, f*) were set to (0.1, 0.2). Heatmaps of estimation accuracy in the 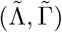 parameter space (tension: a; pressure: d). The black line indicates a boundary, as reported in a previous study [36, 39], that separates two distinct ground state regions. Above this boundary, the ground state is a regular honeycomb network, while below it, the ground states are irregular soft networks characterized by the coexistence of multiple configurations. Gray cells indicate parameter sets where the simulation stopped because of a large distortion of cell morphology. Colors in each cell represent the time average of the correlation coefficient between true and estimated values up to *t* = 30.0. Dependence of estimation accuracy on 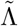 (junctional tension: b; cellular pressure: e) and 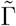 (junctional tension: c; cellular pressure: f).

**Figure 5:**
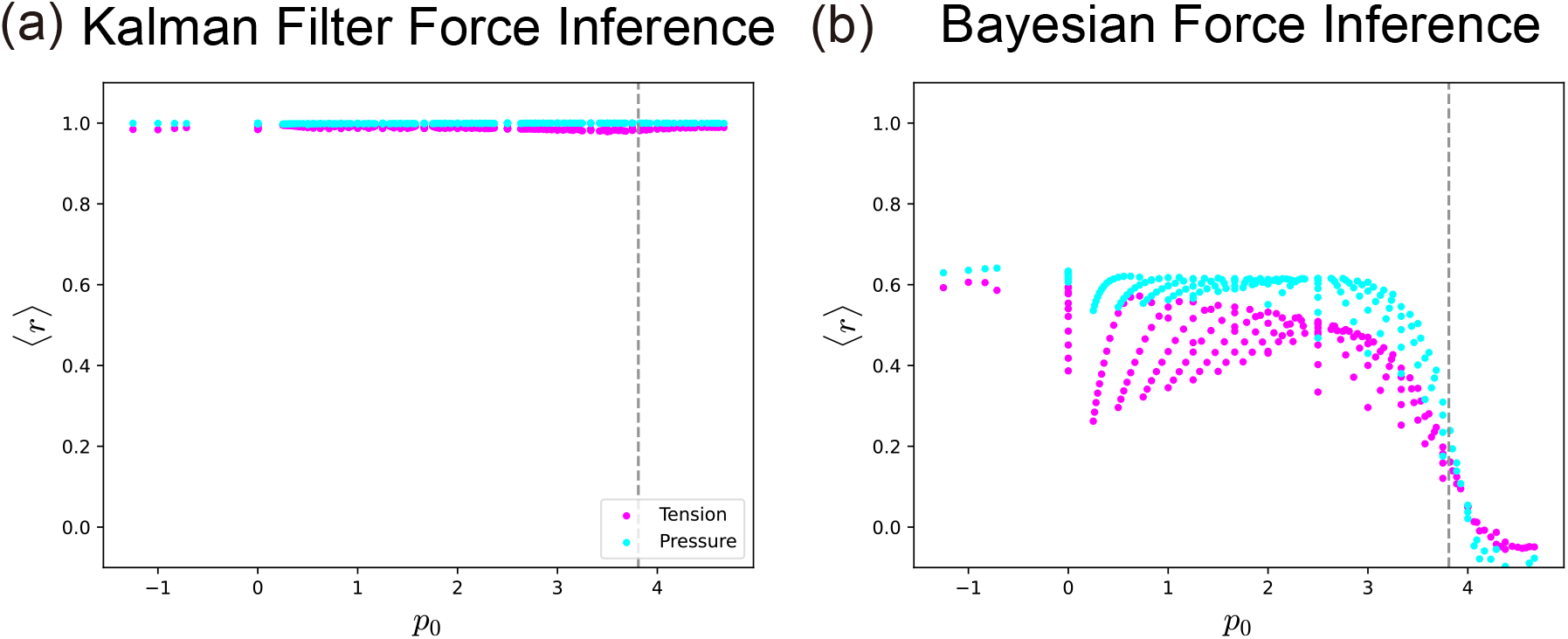
Estimation accuracy of Kalman-filter Force Inference (KFI) and Bayesian Force Inference (BFI) against the target shape index *p*_0_. (a, b) The estimation accuracy of KFI (a) and BFI (b) as a function of the target shape index *p*_0_. Magenta and cyan dots respectively represent the time-average correlation coefficient ⟨*r*⟩ for junctional tension and cellular pressure up to *t* = 30.0. The vertical dashed lines indicate *p*_0_ = 3.81, which is the critical value for solid-fluid transition.

Related to the two regimes, hexagonal and soft-networks, the cell vertex model is known to show a rigidity transition from a solid state to a fluid state, which is governed by a dimensionless parameter target called shape index 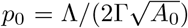 [40]. We found that KFI, as shown above, exhibited high estimation accuracy independently of *p*_0_ [Fig.5 (a)], whereas the estimation accuracy of BFI began to decline around *p*_0_ = 3.0, close to the solid-to-fluid transition point. In the fluid-state, the accuracy was dropped below 0.2 [Fig.5 (b)]. In summary, Kalman-filter Force Inference accurately estimated cellular force across a wide range of tissue material properties, owing to its use of a prior distribution with minimum assumptions.

### 2.4 The robustness of KFI to observation error

So far, we have not considered the influence of noise in the cell vertex position that arise during the measurement process. In practice, vertex positions extracted from the microscopy data inevitably include noise arising from the image sensor pixelation and image analysis (e.g. segmentation), which can affect force inference. A key advantage of the Kalman filter is its ability to provide error estimates for the inferred states by weighing the likelihood function and the prior distribution via the Kalman gain (Eq. 15), thereby contributing to robust estimation even in the presence of noise in the observations [29].

Here, we assessed the tolerance of KFI for such measurement noise using synthetic data with noise. For this, we added Gaussian noise with zero mean and variance 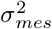 to the vertex coordinates in the synthetic data used in Fig. 2 (a). For the noise strength *σ*_*mes*_, we first considered the typical variation ⟨Δ*R*⟩ in the vertex coordinates induced by the external forcing. The average variation ⟨Δ*R*⟩ is obtained by Eq. 24 in the Materials and Methods. Then, *σ*_*mes*_ was set to 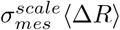, where 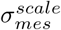 is the scaled noise strength. The observation noise *σ*_*obs*_ in Eq. 10 was set to be 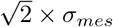. This is because KFI uses displacement (the difference of vertex coordinates) as observation data, whose observation error variance is thus twice that of the vertex coordinates due to the error propagation.

Figs. 6 (a-c) present the estimation results when KFI is applied to the data with 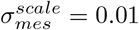. The spatial distribution of the estimated junctional tensions and cellular pressures is quite similar to that in Fig. 2 (b) [Fig. 6 (a)]. While the correlation between the ground truth and the KFI estimates was slightly more scattered than that in Fig. 2 (c), the correlation coefficient remained sufficiently high [Fig. 6 (b)]. We then analyzed the estimated force dynamics of a specific junction and cell [Fig. 6 (c)]. After the initial transient response with undershoot and overshoot, the estimation followed the true values with subtle fluctuations. The 95% HDI from the noisy data was wider than that in Fig.2 (e), reflecting the uncertainty arising from observation noise. The 95% HDI was considerably wide in the initial response, but subsequently converged, containing the true value.

**Figure 6:**
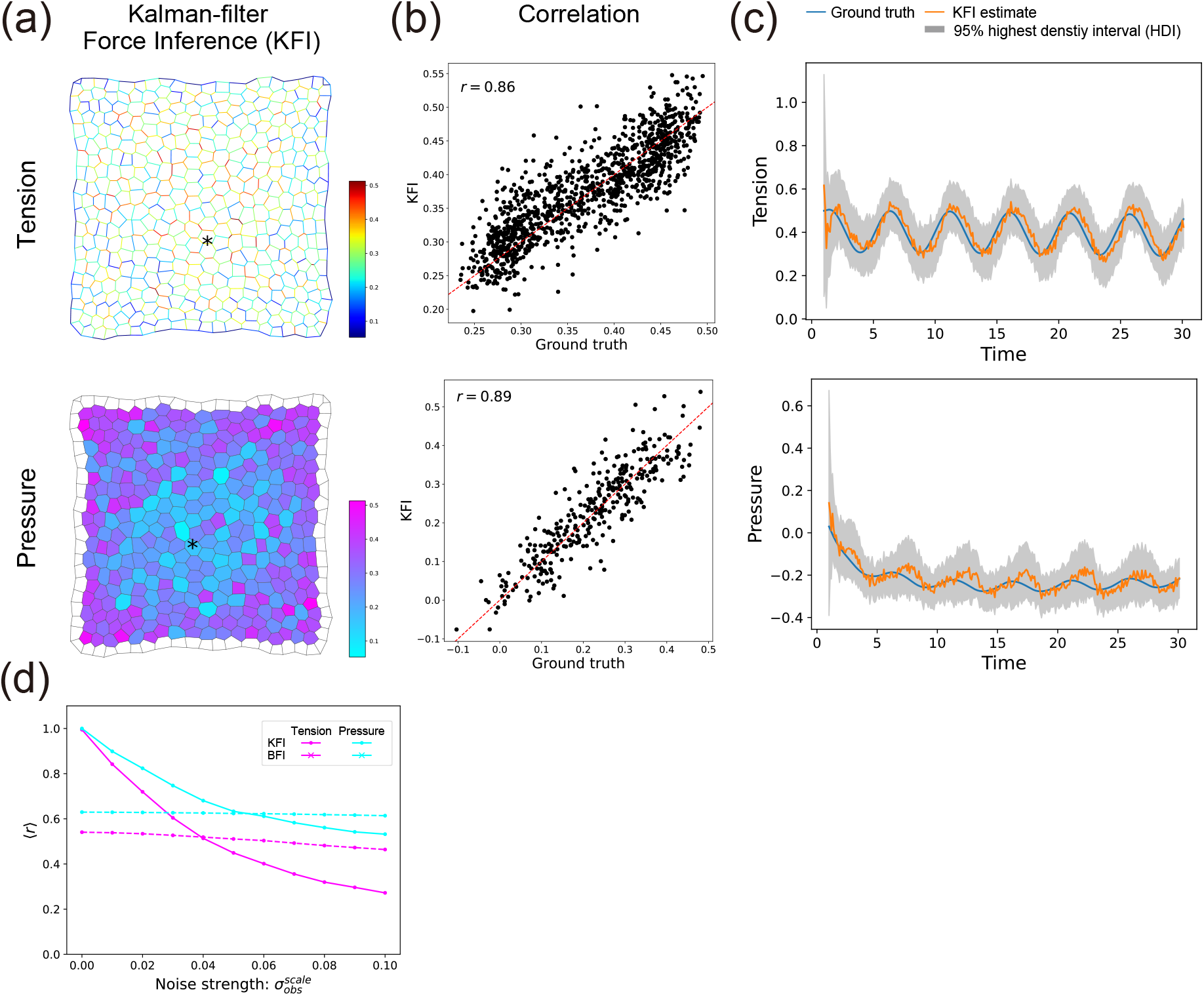
Robustness of Kalman-filter Force Inference (KFI) to observation noise. (a-c) KFI estimation from noisy observation data. Noisy data were generated by adding Gaussian noise to x- and y-coordinates of vertices to the data used in Fig. 2. In panels (a-c), upper and lower plots shows the results for junctional tension and cellular pressure, respectively. (a) The spatial distribution of KFI estimate at *t* = 30.0. The color maps indicate the magnitude of the inferred forces. (b) Correlation between the ground truth and the KFI estimates of the cellular force at *t* = 30.0. The dashed line indicates the *y* = *x* line. The value in the top-left of each panel is the Pearson correlation coefficient. (c) Temporal dynamics of the ground truth and the KFI estimates of cellular forces. The ground truth (blue line) and the KFI estimate (orange line) are plotted against time. The gray shaded area indicates the 95% highest density interval of the estimates. The corresponding junction and cell are indicated by the asterisk in the top and bottom panels of (a), respectively. (d) The time-averaged correlation coefficient, ⟨*r*⟩, as a function of the standard deviation of the observation noise, 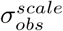. Magenta and cyan lines correspond to junctional tension and cellular pressure, respectively. The lines with dot (•) and cross (×) markers correspond to estimates KFI and Bayesian Force Inference (BFI), respectively.

Next, to study how the estimation accuracy of KFI depends on the strength of measurement noise, we analyzed the time-averaged correlation coefficient ⟨*r*⟩ between the ground truth and the KFI estimate at various strengths of 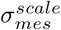. As noise increased, ⟨*r*⟩ decreased, which was more prominent for junctional tension than for cellular pressure [Fig. 6 (d)]. This is because the coefficients for cellular pressure estimation (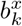 and 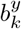 in Eq. 7) are linearly dependent on vertex coordinates, whereas those for junction tension estimation (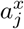 and 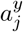 in Eq. 7) exhibit a complex, non-linear dependence on the coordinates. This non-linearity leads to larger errors for tension estimation and a more rapid drop in accuracy. The same analysis performed with BFI revealed that KFI yielded a higher ⟨*r*⟩ than BFI up to 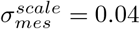 for tension and 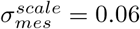 for pressure. Beyond these thresholds, BFI demonstrated superior accuracy. These results suggest that KFI demonstrates adequate robustness to observation noise, while BFI may offer complementary stability, particularly at higher noise levels.

## 3 Discussion

In this study, we introduced the Kalman-filter Force Inference (KFI), a novel method designed to accurately estimate the dynamics of cellular forces from time series data. This method operates within a Bayesian framework that assumes two key principles: namely, the dynamic force balance between cellular and dissipative forces, and the temporal smoothness of cellular forces. This approach naturally leads to an analysis cycle analogous to that of Kalman filter. Our rigorous evaluation, utilizing diverse artificial data generated by the numerical simulations, demonstrated the robust performance of KFI. Specifically, KFI exhibited high estimation accuracy across a wide range of the force dynamics and the tissue material properties, and maintained reasonable accuracy even with errors in the vertex coordinates.

While the force inference methods have significantly advanced, existing approaches often face a fundamental trade-off between robustness and broad applicability. For instance, Bayesian Force Inference (BFI) utilizes a prior distribution for junctional tension, demonstrating significant resilience to observation noise [15, 25, 24]. However, this reliance on a specific prior distribution for tension implies an implicit assumption of underlying cellular mechanical parameters, which restricts their applicability [26]. In contrast, analytical force inference methods, including those for time-lapse movies, typically do not impose such strong prior constraints, making them potentially applicable across a wider range of tissue types. Nevertheless, without explicit prior regularization, these methods can be susceptible to observation noise, leading to estimates that are affected by the noise and therefore exhibit low robustness. In this study, we developed Kalman-filter Force Inference (KFI), a statistical approach that effectively addresses these limitations by leveraging a prior distribution based on the temporal continuity of cellular forces, which enhances robustness against observation noise. Crucially, this prior distribution does not assume specific tension dis-tributions or underlying cellular mechanical parameters, thereby making KFI applicable to tissues with diverse cellular mechanical parameters. By integrating robust noise handling with broad applicability, KFI effectively overcomes the limitations of existing methods.

During the morphogenesis, cell-level mechanical parameters often exhibit heterogeneity in both space and time. Since our method does not rely on a spatiotemporal homogeneity in cell-level mechanical parameters, except for dissipative mechanics, KFI is expected to accurately estimate the cellular forces in tissues with mechanical heterogeneity. We generated a synthetic data with tension that exhibited heterogeneity in time and space [Fig. S5]. When the cortical elasticity Γ gradually increased over time, KFI successfully inferred the cellular forces which also changed over time [Fig. S5 (a-c)]. Even when the line tension exhibited stochastic variations in space and time, it can also accurately estimate the cellular forces [S5 (d, e)]. Thus, our method can be used to unveil a link between spatiotemporal heterogeneity in cell-level mechanical parameters and tissue morphogenesis.

Several improvements are required for the application of KFI to *in vivo* data. For example, the current KFI does not consider changes in cell topology such as cell rearrangement, cell division, and cell death. A straightforward implementation of them involves incorporating cellular forces of all cells and junctions throughout a time-lapse movie into the force vector 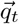, including those that appear or disappear mid-course, following a similar principle to previous work [22].

In the method presented in this paper, the observation noise was considered as an additive noise in Eq. 7 resulting in the likelihood function give by Eq. 10. However, the observation noise in the vertex positions also affect the coefficient matrix *C*_*t*_ in Eq. 7, resulting in a nonlinear influence of the observation noise on the inference process. By considering such effects, the robustness of KFI to observation error will be improved.

Understanding the complex role of dynamic cellular forces in epithelial morphogenesis requires sophisticated analytical methods and KFI can effectively address this need. For example, while the mechanics of collective cell migration have been intensively characterized in cultured cell systems, their *in vivo* characterization remains limited. Furthermore, recent literature highlights that oscillatory behaviors of junctional tension or apical contractile force appear during convergent-extension, and the spatial coordination of these oscillations promotes its efficient progression [41, 42, 43]. Applying KFI to these phenomena would enable accurate visualization of their driving forces, thereby advancing our understanding of the underlying mechanics. It is particularly interesting how the phase pattern in the latter emerges as development progresses. From a physical perspective, relating the cellular forces inferred by KFI to kinematic data, such as the strain rate of junctions and cells, would enable us to extract constitutive equations from time-lapse data, similar to approaches in the previous studies [44, 45]. Moreover, including more detailed friction effects, such as junction friction, is also a worthwhile extension, even though this study primarily considered vertex friction [46, 47, 48, 49, 32]. Through accurate estimation of force dynamics exhibited by cells in tissues with diverse material properties, Kalman-filter Force Inference is expected to contribute to unraveling the mechanisms controlling cellular force dynamics and their spatial patterns in epithelial deformation.

## 4 Materials and Methods

### 4.1 Kalman-filter Force Inference

We implemented the inference algorithm [Fig.2 (a)] using Python. We set the initial values of 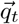 and Σ_*t*_ to be 0.5 and 1.0*I*, respectively. *σ*_*obs*_ was set to be 10^−5^ in sections from 2.2 to 2.3 and 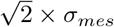 in section 2.4. We set *σ*_*q*_ to be 0.1. For *η* and *dt*, we used their true value in the data generation: *η* = 1.0, *dt* = 0.1.

### 4.2 Bayesian force/stress inference

In this study, we used Bayesian force inference (BFI) as a benchmark [15], available from https://github.com/IshiharaLab/BayesianForceInference. BFI employs a likelihood function derived from the force-balance equation at the cell vertices, equivalent to Eq. 10 with 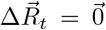. The prior distribution for tension is a normal distribution 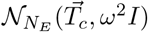, where 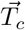 represents a vector of positive tension *T*_*c*_ and the value of *T*_*c*_ is typically set to 1. For pressure, the prior distribution is configured to effectively ignore hydrostatic pressure. These likelihood and prior distributions are combined to compute the posterior distribution, and the marginal likelihood is then numerically maximized with respect to 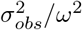 to obtain estimates of cellular forces.

### 4.3 Data generation

In this study, we used synthetic data to assess the accuracy of force inference. The synthetic data on the cell configuration was generated through numerical simulation of the cell vertex model [30, 31, 34]. The positions and connectivity of vertices 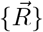 were changed according to the minimization of virtual work 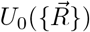. The virtual work is given as

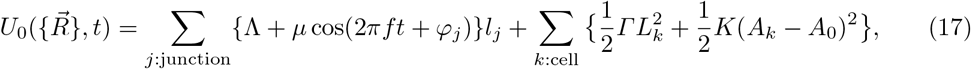

where Λ, *Γ, K* and *A*_0_ are the line tension, cortical elasticity of cell perimeter, area elasticity of a cell and the target area of a cell, respectively. *µ* and *f* are the strength of time dependence and the frequency of the dynamics of the line tension, respectively. *l*_*j*_, *L*_*k*_ and *A*_*k*_ respectively represent the length of the cell-cell junction *j*, the cell perimeter and the cell area of cell *k. φ*_*j*_ is the initial phase of the dynamics of junctional tension. The partial derivatives of the virtual work by *l*_*j*_ and *A*_*k*_ respectively give the junctional tension and cellular pressure:

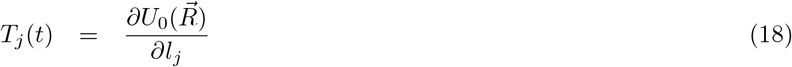

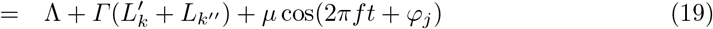

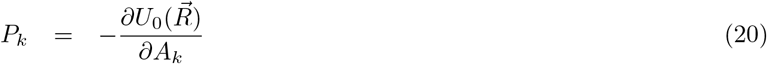

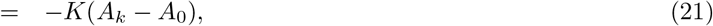

where *k*^*′*^ and *k*^*′′*^ indicate the indices of cells separated by junction j.

We solved the following equation in simulations:

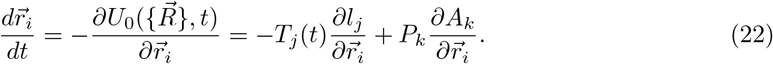

The numerical simulations were implemented by C++. We set an initial cell configuration as a 20 × 20 cell tile, which was generated from randomly distributed centroids with a mean cell area of *A*_0_. *φ*_*ij*_ is initialized with the random number drawn from the uniform distribution from −*π* to *π*. We adopted a free boundary condition.

We generated noisy datasets to assess the robustness of our method. The data presented in Fig.2 were used as the clean base for this process. To define the reference length scale of the observation noise, we first calculated the range of all vertex coordinates during the last period of tension oscillation in the simulation *T*_*last*_. For instance, the range of the x-coordinate of the *i*-th vertex is calculated as:

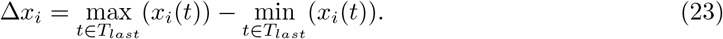

The average range of vertex coordinates was then computed as the reference length scale for the observation noise:

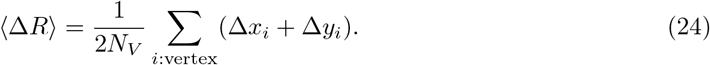

Finally, we generated the noisy data by adding Gaussian noise, 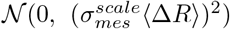, to the x- and y-coordinates of all vertices at each time point of the simulation. We created datasets with varying noise levels by setting 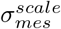 from 0.01 to 0.1 in increments of 0.01.

### 4.4 Measure of accuracy

We used correlation coefficient for measuring estimation accuracy. The correlation coefficient of junctional tension {*T*_*j*_} at time *t* is calculated as below:

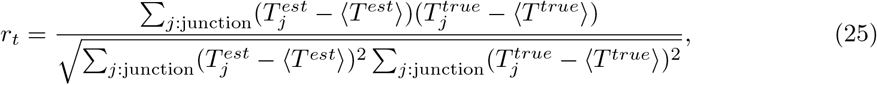

where ⟨*T* ^*est*^⟩ and ⟨*T* ^*true*^⟩ respectively represent the average estimated and true tension at time *t*. The correlation coefficient for the cellular pressure is calculated in the same way as for junctional tension.

## Acknowledgments

We thank K. Sugimura, S. Ishihara, and the members of Laboratory for Physical Biology for their fruitful discussions and helpful comments on the manuscript. The work is part of the RIKEN Pioneering Project “Prediction for Science”. This study was financially supported by JSPS KAKENHI Grant(23K16999) to G.O. and 22H05170 to T.S.. G.O. was supported by RIKEN Special Postdoctoral Researcher Program.

---

**Figure S1:**
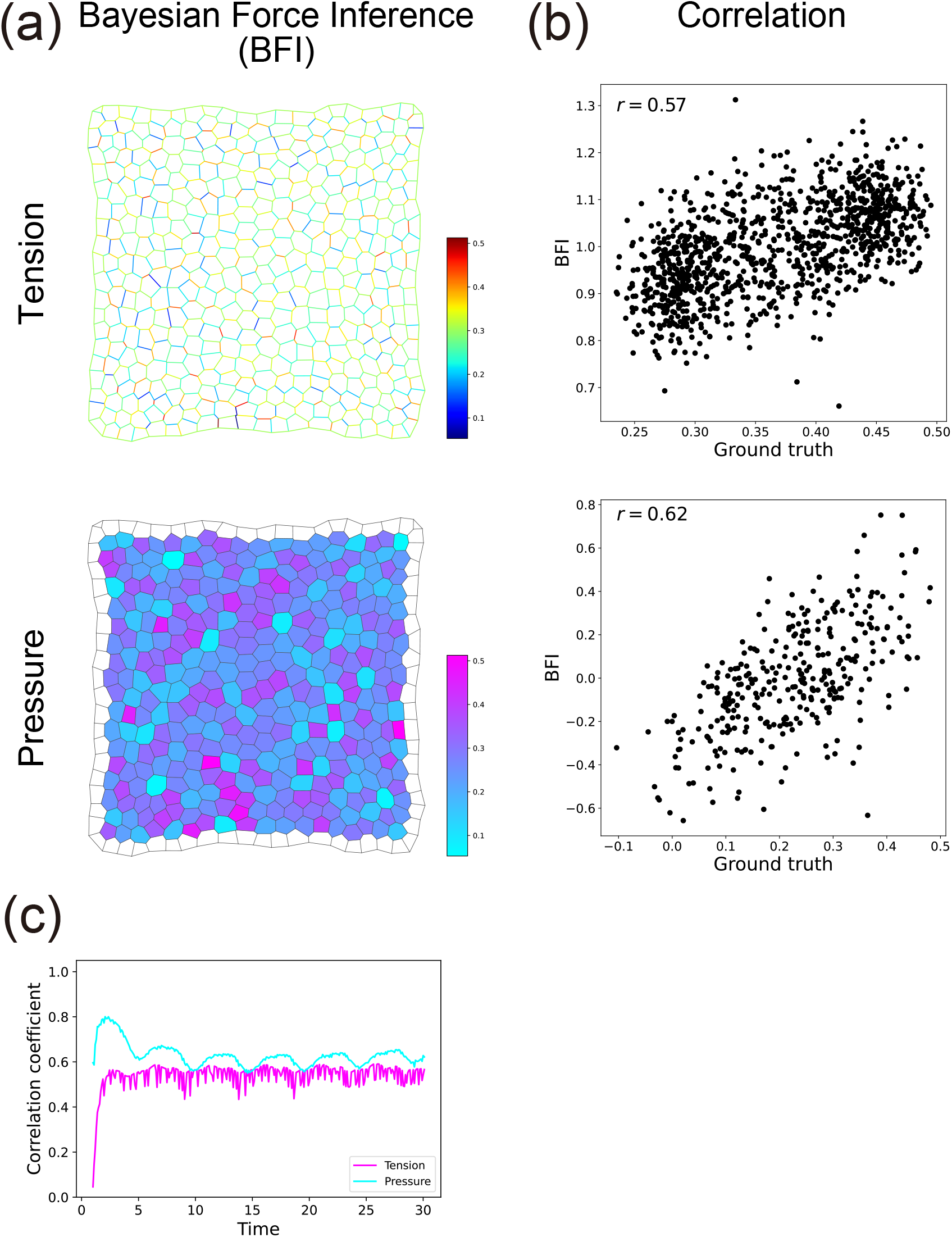
Application examples of Bayesian Force Inference (BFI). (a) The spatial distribution of the ground truth and the BFI estimates of the cellular forces at *t* = 30.0. BFI was applied to the same data in Fig. 2. The top and bottom panels correspond to junctional tensions and cellular pressures, respectively. The magnitude of junctional tension and cellular pressure is indicated by the colors exemplified in the color bar. (b) The correlation between the ground truth and the BFI estimates of the cellular force at *t* = 30.0. The top and bottom panels correspond to junctional tensions and cellular pressures, respectively. The number in the upper left corner of each panel represents the Pearson correlation coefficient between the ground truth and the estimated value by BFI. (c) Correlation coefficients between the ground truth and the BFI estimates over the entire time course. The magenta and cyan lines correspond to junctional tension and cellualr pressure, respectively.

**Figure S2:**
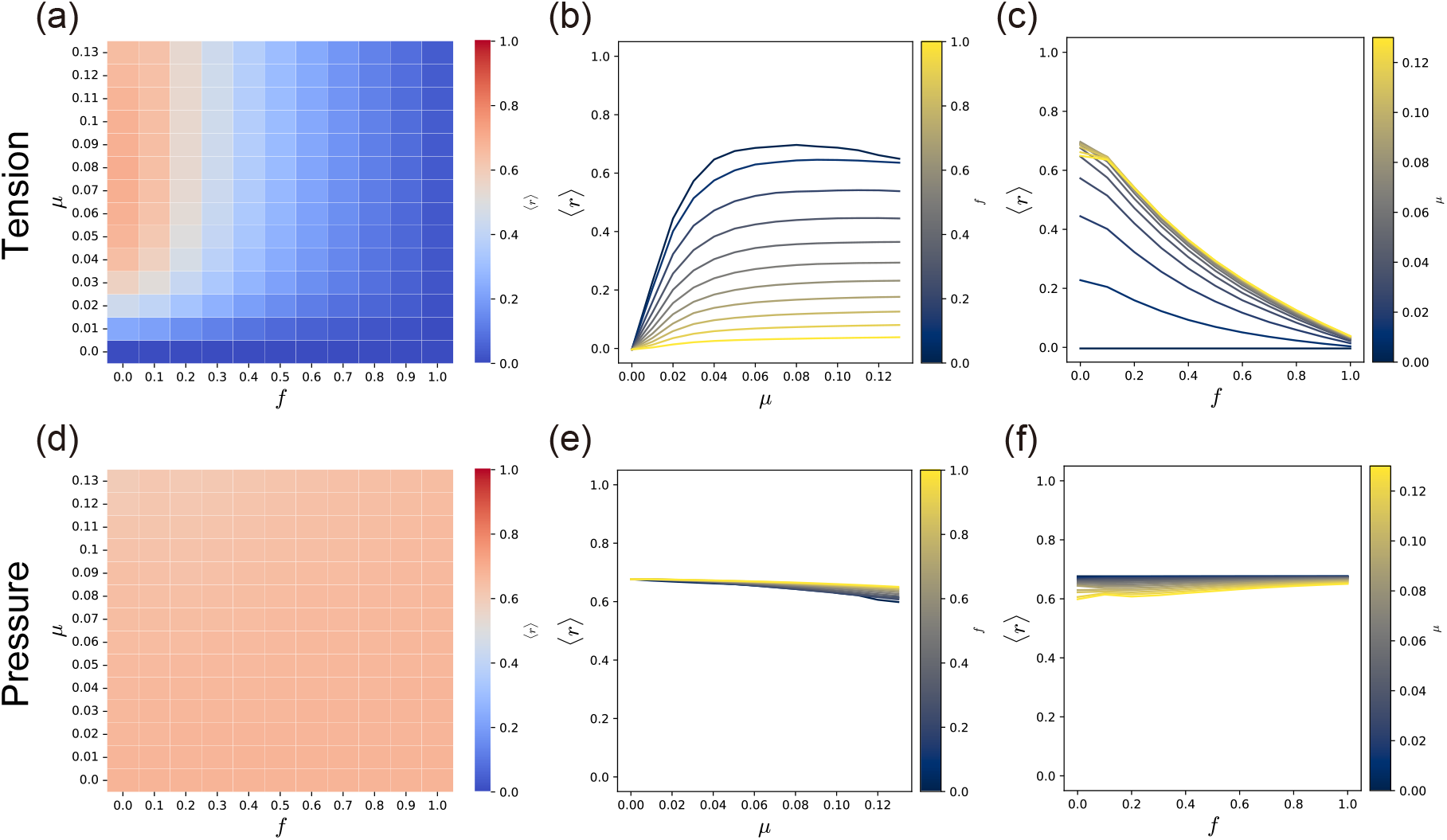
Estimation accuracy of Bayesian force inference (BFI) for various dynamic parameters (*µ, f*). (a-f) Parameter (*µ, f*) dependence of BFI’s estimation accuracy for junctional tension (a-c) and cellular pressure (d-f). Heatmaps of estimation accuracy in the (*µ, f*) parameter space (junctional tension: a; cellular pressure: d). Colors in each cell represent the time average of the correlation coefficient between true and estimated values up to *t* = 30.0. Dependence of estimation accuracy on *µ* (junctional tension: b; cellular pressure: e) and *f* (junctional tension: c; cellular pressure: f).

**Figure S3:**
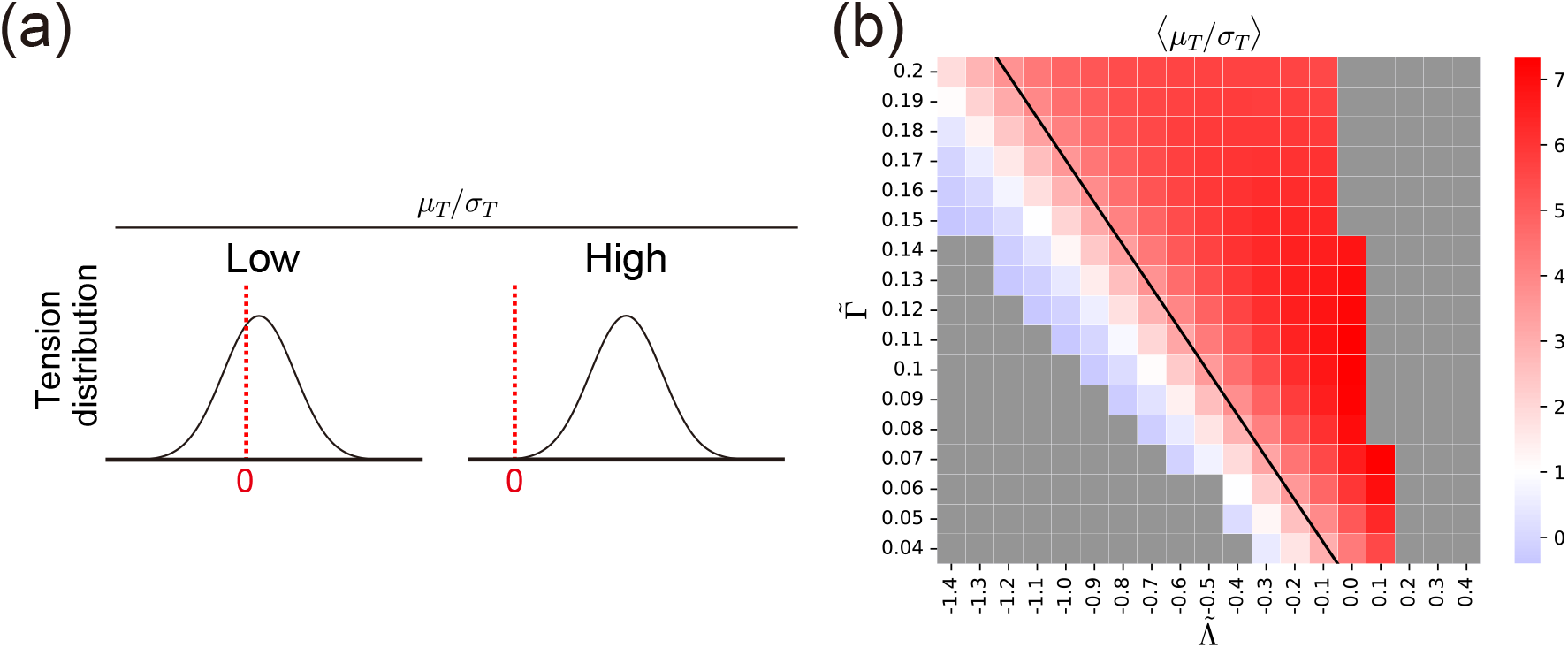
Dependence of tension distribution on material parameters 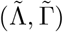. (a) Relationship between junctional tension distribution and the ratio of the mean to the standard deviation of junctional tension *µ*_*T*_ */σ*_*T*_. When *µ*_*T*_ */σ*_*T*_ is small, junctional tension values are broadly distributed around zero, rendering the positive-tension prior of Bayesian Force Inference (BFI) inappropriate (left panel). Conversely, a large *µ*_*T*_ */σ*_*T*_ indicates that most junctional tension values are positive, aligning well with the prior distribution of BFI. (b) Heatmap of the time-averaged *µ*_*T*_ */σ*_*T*_ in the 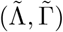 parameter space.

**Figure S4:**
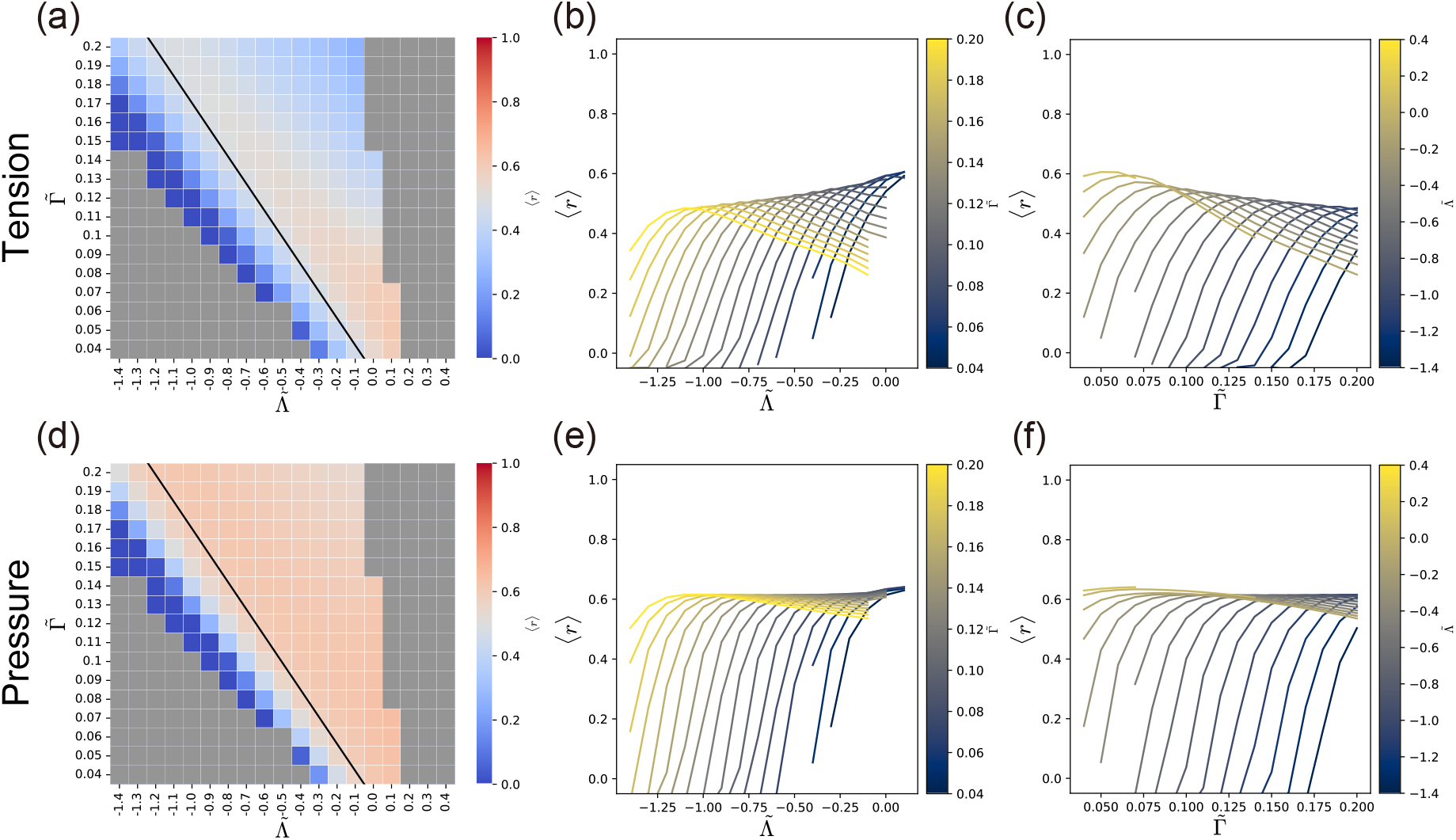
Estimation accuracy of Bayesian force inference (BFI) for various mechanical parameters 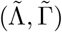. (a-f) Parameter 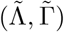 dependence of BFI’s estimation accuracy for junctional tension (a-c) and cellular pressure (d-f). The dynamics parameters (*µ, f*) were set to (0.1, 0.2). Heatmaps of estimation accuracy in the 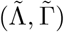 parameter space (junctional tension: a; cellular pressure: d). The black line represents the boundary, as explained in the caption of Fig. 4. Gray cells indicate parameter sets where the simulation stopped because of a large distortion of cell morphology. Colors in each cell represent the time average of the correlation coefficient between true and estimated values. Dependence of estimation accuracy on 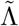 (junctional tension: b; cellular pressure: e) and 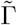 (junctional tension: c; cellular pressure: f).

**Figure S5:**
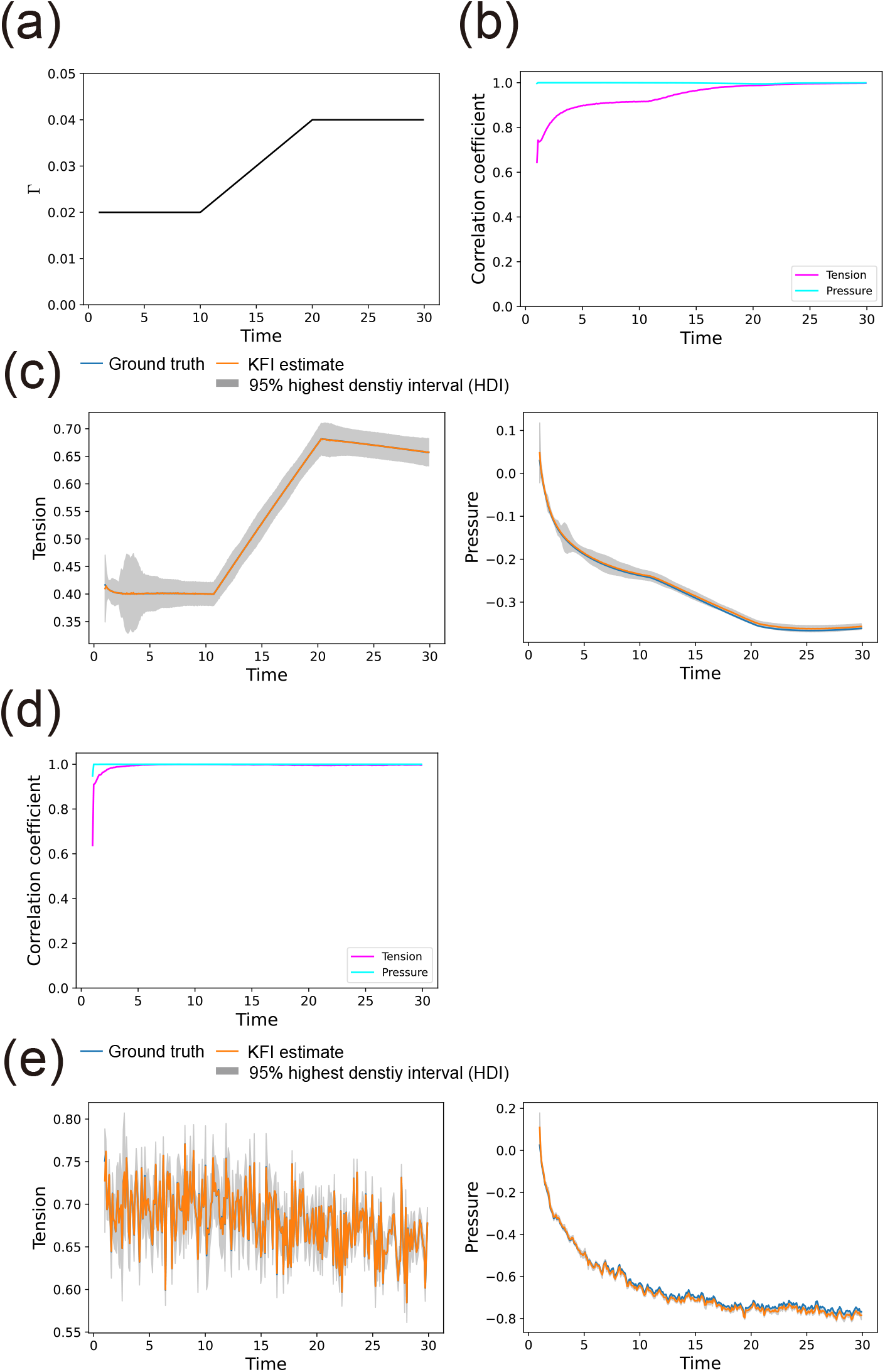
Estimation accuracy of Kalman-filter Force Inference (KFI) for various dynamic forces. (a-c) Time-varying mechanical parameter. The true tension was calculated by the conventional model [36, 37]: 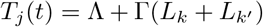. (a) The dynamics of mechanical parameter Γ. Γ was set to 0.02 (*t* ≤ 10), linearly increased to 0.04 (10 *< t* ≤ 20), and held constant at 0.04 (*t >* 20). Other parameters were set as Λ = 0.1, and *k* = *A*_0_ = 1.0. (b) Correlation coefficients between the ground truth and the KFI estimates over the entire time course. The magenta and cyan lines correspond to junctional tension and cellular pressure, respectively. (c) Temporal dynamics of the ground truth and the KFI estimates of cellular forces (left: junctional tension; right: cellular pressure). The ground truth (blue line) and the estimate (orange line) are plotted against time. The gray shaded area indicates the 95% highest density interval (HDI) of the estimates. (d-e) Stochastic junctional tension. The true junctional tension was calculated by 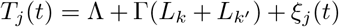. The noise *ξ*_*j*_(*t*) is a correlated Gaussian noise with ⟨*ξ*_*j*_(*t*)⟩ = 0 and ⟨*ξ*_*j*_(*τ*)*ξ*_*j*_(0)⟩ = 0.05 exp(−0.1*τ*), where *τ* =| *t*| represents the time lag. Other parameters were set as (Λ, Γ, *k, A*_0_) = (0.1, 0.04, 1.0, 1.0). (d) Correlation coefficients between the ground truth and the KFI estimates over the entire time course. (e) Temporal dynamics of the ground truth and the KFI estimates of cellular forces (left: junctional tension; right: cellular pressure).

